# miR-122 affects both the initiation and maintenance of Hepatitis C Virus infections

**DOI:** 10.1101/2021.11.04.467384

**Authors:** Mamata Panigrahi, Patricia A Thibault, Joyce A Wilson

## Abstract

A liver-specific microRNA, miR-122, anneals to the HCV genomic 5’ terminus and is essential for virus replication in cell culture. However, bicistronic HCV replicons and full length RNAs with specific mutations in the 5’ UTR can replicate, albeit to low levels, without miR-122. In this study, we have identified that HCV RNAs lacking the structural gene region or having EMCV IRES-regulated translation had reduced requirements for miR-122. In addition, we found that a smaller proportion of cells supported miR-122-independent replication when compared a population of cells supporting miR-122-dependent replication, while viral protein levels per positive cell were similar. Further, the proportion of cells supporting miR-122-independent replication increased with the amount of viral RNA delivered, suggesting that establishment of miR-122-independent replication in a cell is affected by amount of viral RNA delivered. HCV RNAs replicating independent of miR-122 were not affected by supplementation with miR-122, suggesting that miR-122 is not essential for maintenance of a miR-122-independent HCV infection. However, miR-122 supplementation had a small positive impact on miR-122-dependent replication suggesting a minor role in enhancing ongoing virus RNA accumulation. We suggest that miR-122 functions primarily to initiate an HCV infection but has a minor influence on its maintenance, and we present a model in which miR-122 is required for replication complex formation at the beginning of an infection, and also supports new replication complex formation during ongoing infection and after infected cell division.

**IMPORTANCE:** The mechanism by which miR-122 promotes the HCV life cycle is not well understood, and a role in directly promoting genome amplification is still debated. In this study, we have shown that miR-122 increases the rate of viral RNA accumulation and promotes the establishment of an HCV infection in a greater number of cells than in the absence of miR-122. However, we also confirm a minor role in promoting ongoing virus replication and propose a role in the initiation of new replication complexes throughout a virus infection. This study has implications for the use of anti-miR-122 as potential HCV therapy.

## INTRODUCTION

Hepatitis C virus (HCV) is a positive strand RNA flavivirus that primarily infects the liver and is carried by over 71 million people worldwide (1). Around 70% of infections cause chronic Hepatitis C disease, which can lead to complications such as liver cirrhosis, hepatocellular carcinoma, and decompensated liver disease (2). Chronic HCV was the primary cause of liver transplantations in the United States, Europe, and Japan for many years, but can now be cured using highly effective direct acting antiviral (DAA) treatments (3).

The positive strand RNA genome of HCV is approximately 9.6 kb long and contains a single open reading frame (ORF) that encodes a viral polyprotein. The ORF is flanked by 5’ and 3’ untranslated regions (UTR) that have secondary structures essential for viral translation and replication (4, 5). The uncapped HCV 5’UTR bears an internal ribosomal entry site (IRES) that directs cap-independent translation of the viral polyprotein, which is subsequently cleaved by viral and cellular proteases into three structural (core, E1, and E2) and seven non-structural (p7, NS2, NS3, NS4a and 4b, NS5a and 5b) viral proteins. The non-structural proteins NS3-NS5b form the replicase complex and are the minimum viral proteins required for genome replication (6, 7).

Micro RNAs (miRNA) are small non-coding RNAs (approximately 22 nucleotides) that regulate gene expression by suppressing mRNA translation and promoting mRNA decay (8). However, contrary to the conventional suppressive role to miRNAs, the liver-specific miRNA, miR-122, anneals to the HCV genome and is essential for viral propagation (9). miR-122, in association with human Argonaute proteins (Ago 1-4) (hAgo:miR-122 complex), binds to two sites on the extreme 5’UTR of the viral genome and promotes viral RNA accumulation (**Fig. 1**) (9–11). While the precise mechanism of HCV replication promotion remains incompletely understood, different roles of miR-122 have been identified, including translation stimulation (12– 14), genome stabilization (15–17), and a direct role in genome amplification (18). Recent studies have also predicted that miR-122 annealing to the viral 5’UTR shapes the structure of the viral 5’UTR RNA and promotes the formation of an “active” or a translation favourable viral IRES structure (14, 19–21); however, biophysical analyses are still required to support this hypothesis.

**FIG 1:**
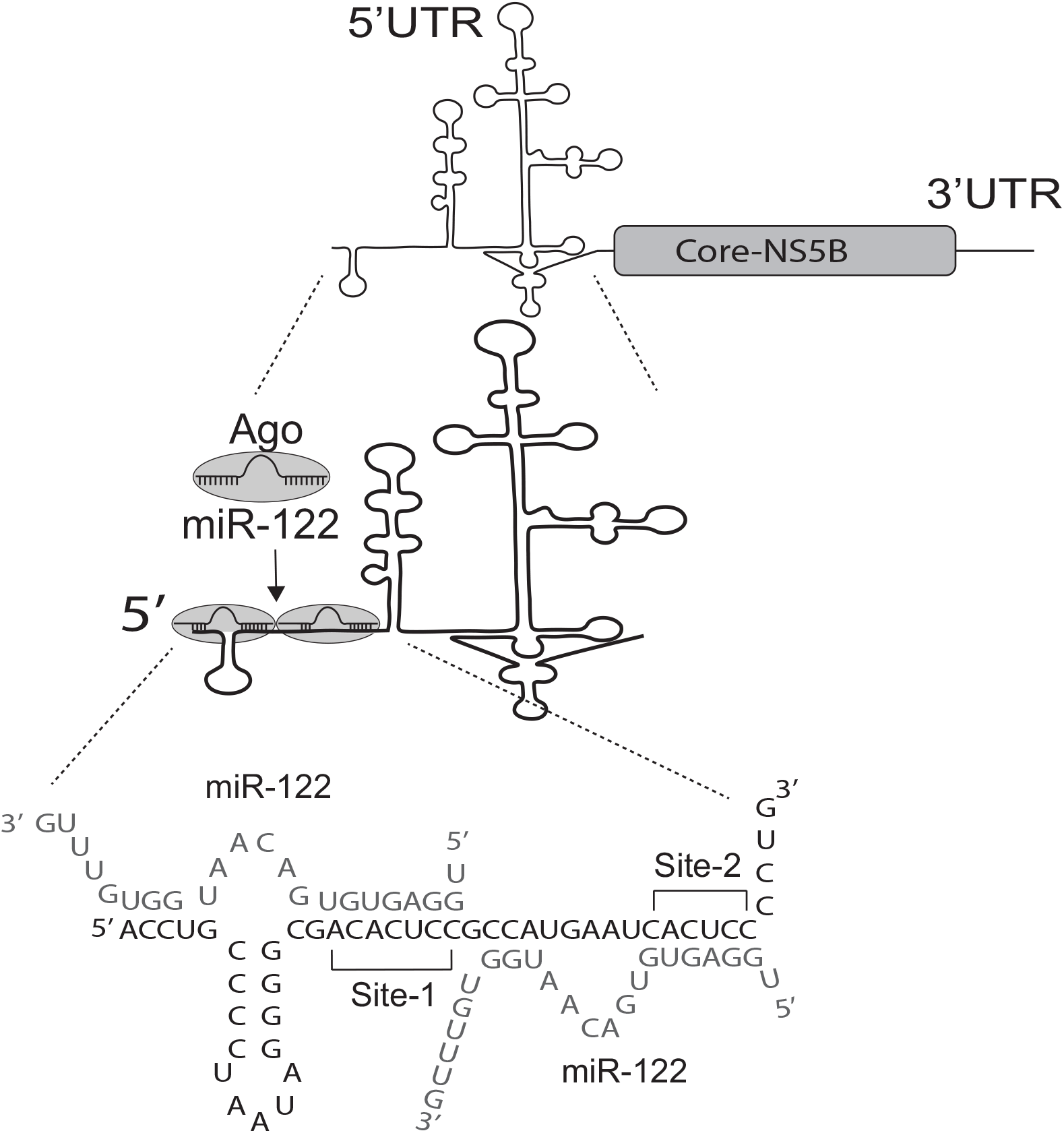
miR-122 annealing to the HCV genome. An Ago:miR-122 complex binds to two sites (Site1 and Site2) on the 5’UTR of HCV genome and promote viral propagation. The miR-122 binding pattern and sequence of 1-42 nucleotide of the 5’UTR of Hepatitis C Virus genome and miR122 annealing to it shown.

Since miR-122 is an important host factor for HCV replication, miR-122 antagonists have shown promising results in a prolonged reduction of viral load in both preclinical and clinical trials (22–24), but the recent emergence of resistant HCV mutants (25) suggests that the virus can adapt to grow in the absence of miR-122. In addition, lab-derived HCV mutants capable of low-level miR-122-independent replication have been identified. These viruses are either bicistronic, with virus protein translation mediated by an EMCV IRES, or have specific mutations in the miR-122 binding sites of the 5’ UTR (19, 20, 25–30). By contrast, wild-type genome replication is undetectable in the absence of miR-122, even with the use of sensitive reporter genomes. Since miR-122 expression is specific to the liver and comprises almost 70% of the total small RNA of the liver (31), it likely plays a role in regulating HCV’s liver tropism. However, there have been reports of extrahepatic HCV RNA and replication (2, 32, 33) and chronic HCV is associated with a broad range of extrahepatic conditions such as cryoglobulinemic vasculitis, diabetes mellitus, and B-cell lymphoma, suggesting a possible role for miR-122-indepenedent HCV replication in these manifestations (2, 34). Further, others have shown selection of cells supporting stable miR-122-independent replication of HCV replicon RNA that suggested a role of miR-122 for establishment, but not maintenance of replication (25).

In this study, we aimed to compare transient and stable miR-122-dependent *vs*. miR-122-independent HCV replication to identify HCV genetic elements that modulate HCV’s dependency on miR-122 and determine the roles of miR-122 in different stages of the HCV life cycle. We have found that the presence of an EMCV IRES and a smaller genome size allows HCV to replicate independent from miR-122, suggesting roles for modified translation regulation and smaller genome sizes in facilitating miR-122-independent replication. Using several models of miR-122-independent HCV replication we have also found that miR-122-independent replication appears efficient within individual cells but is established in only a small number of cells in the population. Further, we also found that miR-122 has a dramatic impact during the early stages and the establishment of an HCV infection, but a less potent supportive role for ongoing genome amplification. From this, we propose a mechanistic model where miR-122 promotes translation and genomic RNA stability early in the virus life cycle to ‘jump-start’ a virus infection, but also has a lesser role in maintaining an ongoing infection, which we propose is to initiate new virus replication complexes as the infection amplifies within a cell and as liver cells divide.

## RESULTS

### EMCV IRES-mediated translation of the HCV non-structural genes promotes miR-122-independent HCV replication

In previous works, we have identified that the HCV subgenomic replicon SGR JFH-1 Fluc was capable of genome amplification independent of miR-122 in Hep3B cells (26). This construct lacks the structural proteins core, E1, E2, and p2, as well as non-structural protein NS2; while translation of the Fluc reporter gene is driven by the HCV endogenous IRES, translation of the non-structural polyprotein (NS3 through NS5b) is driven by an EMCV IRES. These data suggested that one or all of; a) the addition of an EMCV IRES to regulate viral non-structural gene translation, b) the removal of the structural proteins, or c) the reduced genome length permitted HCV SGR RNA replication independent of miR-122. In this study, we aimed to determine the independent influence of the EMCV IRES, the structural gene region, and the size of the RNA genome on miR-122 independent replication.

To determine if the addition of an EMCV IRES facilitated miR-122-independent replication, we assessed transient miR-122-dependent and miR-122-independent replication of full-length HCV RNAs in which viral gene translation was regulated by an EMCV IRES (Fig. 2A) *i*.*e* FGR JFH-1 Fluc which has a firefly luciferase reporter gene, or J6/JFH-1 Neo (p7Rluc2a) which has a *Renilla* luciferase reporter gene and a Neomycin selection marker (herein noted as J6/JFH-1 Neo Rluc). To assess miR-122-independent and miR-122-dependent replication, the viral RNAs were co-electroporated into Huh 7.5 miR-122 knock out (KO) cells with either control microRNA (miControl) to assess miR-122-independent replication, or synthetic miR-122 to assess miR-122-dependent replication (Fig. 2B-E). HCV genomic RNA amplification was assessed based on luciferase expression as a proxy for virus replication and SGR JFH-1 Fluc RNA was included as a positive control for miR-122-independent replication (26). All of the viral RNAs replicated efficiently when miR-122 was added **(Fig. 2B-E**) and as we and others have consistently observed, the J6/JFH-1 (p7Rluc2a) full-length wild-type construct (herein noted as J6/JFH-1 Rluc) does not display miR-122-independent HCV replication. However, both the bicistronic full-length genomic replicon expressing a firefly luciferase reporter gene (FGR JFH-1 Fluc) and a bicistronic full-length genomic replicon that expresses both a neomycin selection marker and a *Renilla* luciferase reporter gene (J6/JFH-1 Neo Rluc) (**Fig. 2A**) showed miR-122-independent replication since they demonstrated measurable luciferase levels above their corresponding GNN or GND negative controls (**Fig. 2B-F**). These results suggest that translation facilitation by the EMCV IRES allows both full-length and sub-genomic HCV to replicate in the absence of miR-122 and that translation regulation of the viral proteins by the EMCV IRES can partially compensate for the lack of miR-122. This experiment additionally rules out the contribution of a given reporter gene, Fluc or Rluc, to miR-122-independent replication (**Fig. 2 C, D, E)**. Thus, altered HCV translation regulation by the addition of the EMCV IRES facilitates virus replication in the absence of miR-122. However, miR-122-independent replication of SGR JFH-1 Fluc, which lacks the structural gene region, is more efficient than that of FGR JFH-1 Fluc, in which has the structural gene region is intact (**Fig. 2F**) suggesting that the structural gene region may inhibit, or shorter genome length enhance miR-122-independent replication of HCV.

**FIG 2:**
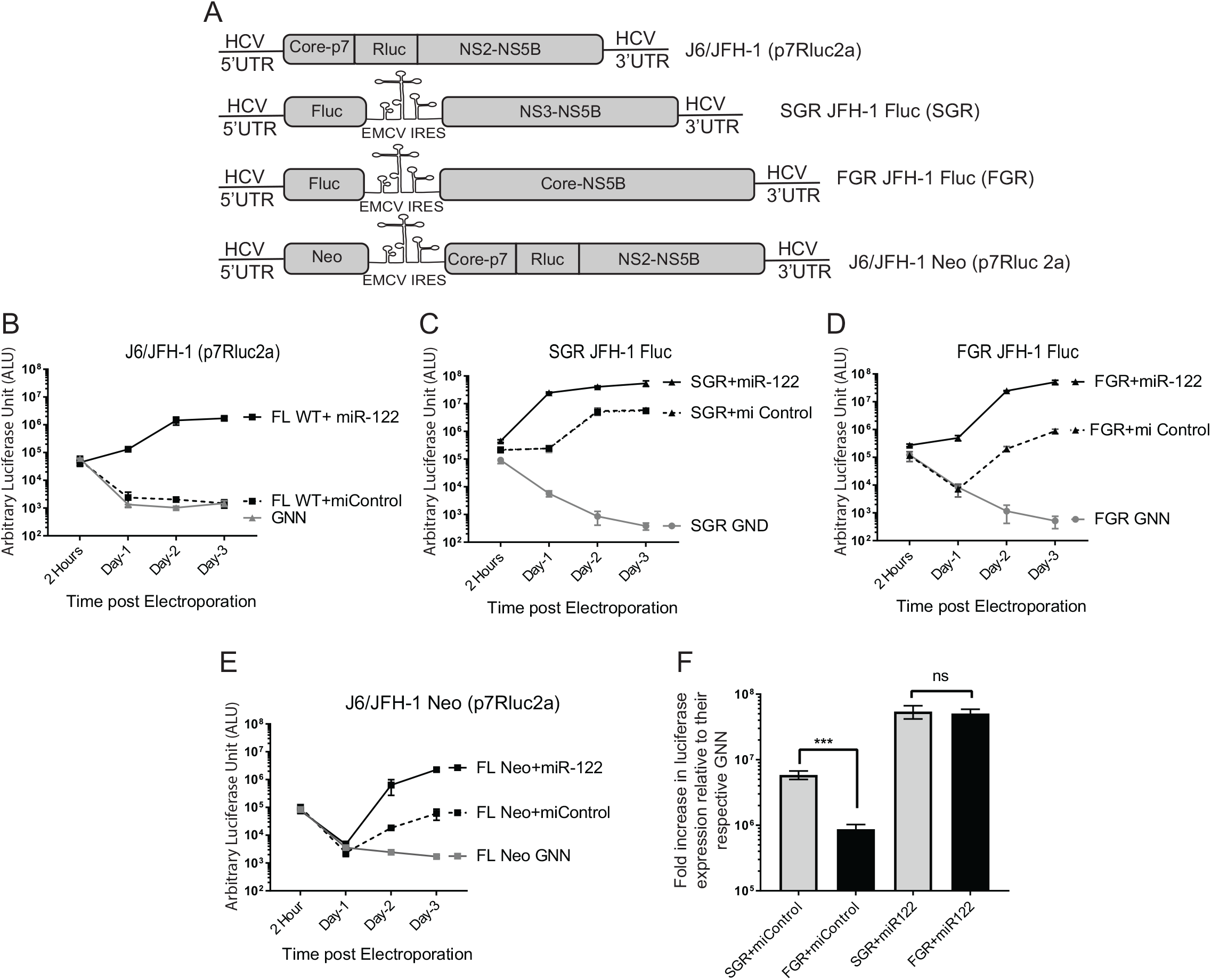
HCV RNAs containing an EMCV IRES replicate independent from miR-122. (A) Schematic diagram of the HCV viral genomes. Top to bottom: Monocistronic Full-Length wild-type J6/JFH-1 (p7Rluc2a) with a *Renilla* luciferase reporter, Bicistronic sub-genomic Replicon (SGR) JFH-1 with a Firefly luciferase reporter SGR JFH-1 Fluc, Bicistronic Full-Length genomic replicon (FGR) or FGR JFH-1Fluc with a Firefly luciferase reporter, Bicistronic Full-Length wild-type J6/JFH-1 Neo (p7Rluc2a) with a *Renilla* luciferase reporter and a Neomycin selection marker. (B, C, D, E, and F) Transient replication of HCV viral genomes shown in (A) in the presence of a control miRNA (miControl) or miR-122 in Huh 7.5 miR-122 knockout cells measured by luciferase reporter expression. (B) J6/JFH-1 (p7Rluc2a), (C) SGR JFH-1 Fluc, (D) FGR JFH-1 Fluc and (E) J6/JFH-1 Neo (p7Rluc2a). Replication defective version of each genome, GNN or GND are negative controls. (F) Fold change in luciferase expression vs non-replicative GNN controls of a full length (FGR JFH-1 Fluc) and a subgenomic (SGR JFH-1 Fluc) HCV adapted from data shown in (B) and (C). All data shown are the average of three or more independent experiments and error bar indicates the standard deviation of the mean. The significance was determined by using Student’s t-test (ns-not significant, ***P<0.001).

### Shorter genomic RNAs support miR-122-independent HCV replication

To determine the contribution of the structural gene region and the size of the RNA genome to miR-122-independent replication, we assessed transient miR-122-dependent and miR-122-independent replication of HCV RNAs in which various regions, or all of the structural gene region was removed. For this we designed several monocistronic constructs derived from J6/JFH-1 (p7Rluc2a) (lacking the EMCV IRES) where we have deleted part, or all, of the structural gene region (**Fig. 3A**). The deletion mutants were designed such that they did not adversely affect polyprotein processing, and all were replication competent in the presence of miR-122 **(Fig. 3B-F)**. However, only two mutants in which the entire structural gene region was removed, SGR JFH-1 (Rluc2a) NS2, and SGR JFH-1 (Rluc2a) NS3 were capable of replicating in the absence of miR-122 (**Fig. 4 A, B, and C**). Thus, we concluded that the absence of the structural genes in their entirety, rather than a specific protein coding region, modulated the ability of HCV RNA to replicate independent from miR-122, and we speculate that may be due to a general increase in rate of genomic RNA replication based on a smaller genome size.

**FIG 3:**
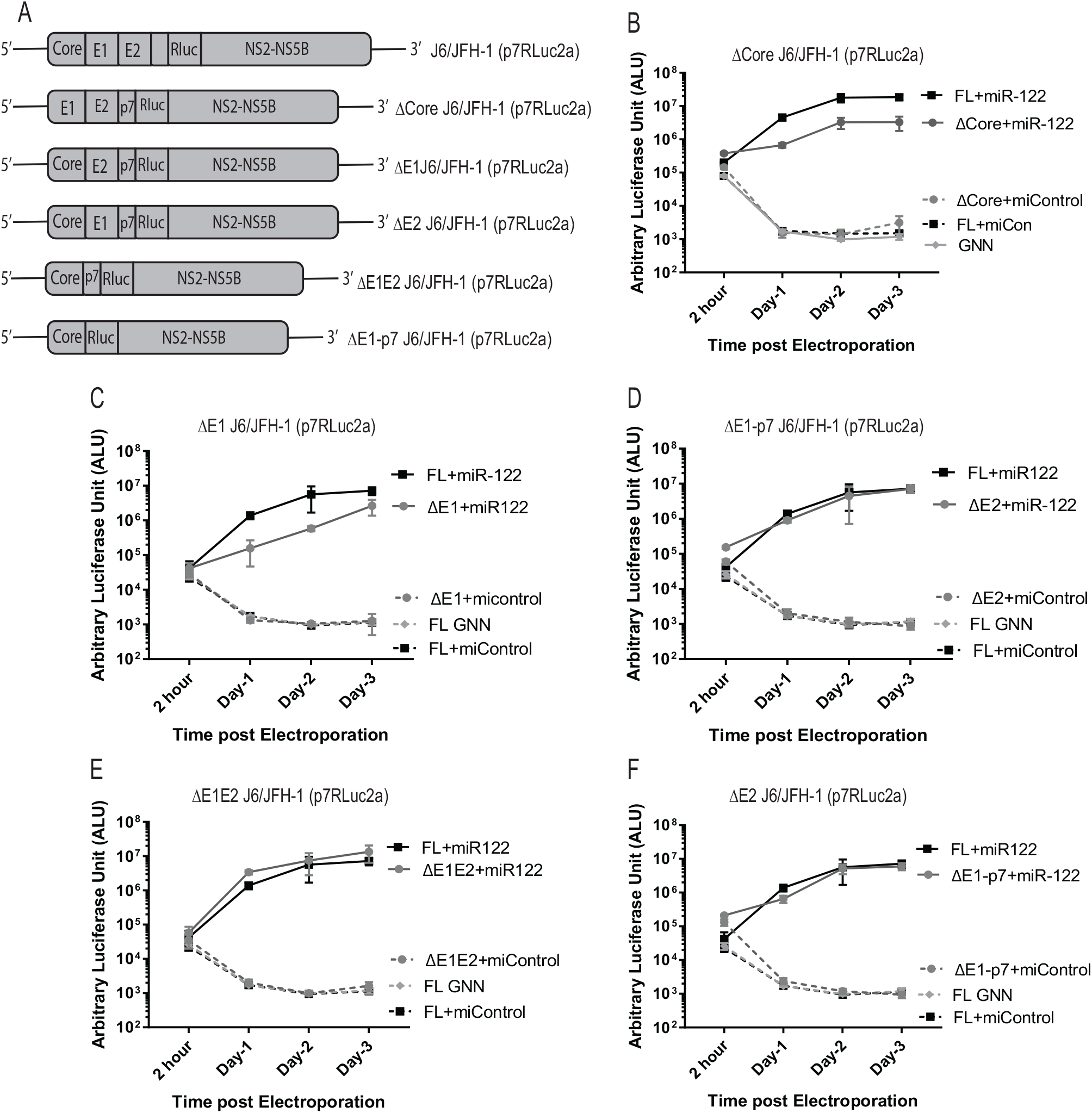
Deletion of the portions of the structural gene region does not facilitates miR-122 independent replication of HCV. (A) Schematic diagrams of the structural gene deletion mutant HCV constructs used, from top to bottom: full length WT, J6/JFH-1(p7Rluc2a); J6/JFH-1 with deletion of core region, Δcore J6/JFH-1 (p7Rluc2a); J6/JFH-1 with deletion of E1 region, ΔE1 J6/JFH-1 (p7Rluc2a), J6/JFH-1 with deletion of E2 region, ΔE2 J6/JFH-1 (p7Rluc2a), J6/JFH-1 with deletion of E1E2 region,ΔE1E2 J6/JFH-1 (p7Rluc2a); and J6/JFH-1 with deletion of E1-p7 region, ΔE1-p7 J6/JFH-1 (Rluc2a). Transient replication of structural gene deletion HCV RNAs in Huh 7.5 miR-122 KO cells in presence of either control microRNA (miControl) or miR-122. (B) WT J6/JFH-1 Rluc HCV RNA and Δcore J6/JFH-1 (p7Rluc2a) RNA; (C) ΔE1 J6/JFH-1 (p7Rluc2a) RNA; (D) ΔE2 J6/JFH-1 (p7Rluc2a) RNA; (E) ΔE1E2 J6/JFH-1 (p7Rluc2a) RNA; and (F) ΔE1-p7 J6/JFH-1 (Rluc2a). A replication defective GNN mutant was included for each RNA as a negative control. All the experiments are a representation of 3 or more replicates. The significance was determined by using Student’s t-test (ns-not significant, *P<0.033).

**FIG 4:**
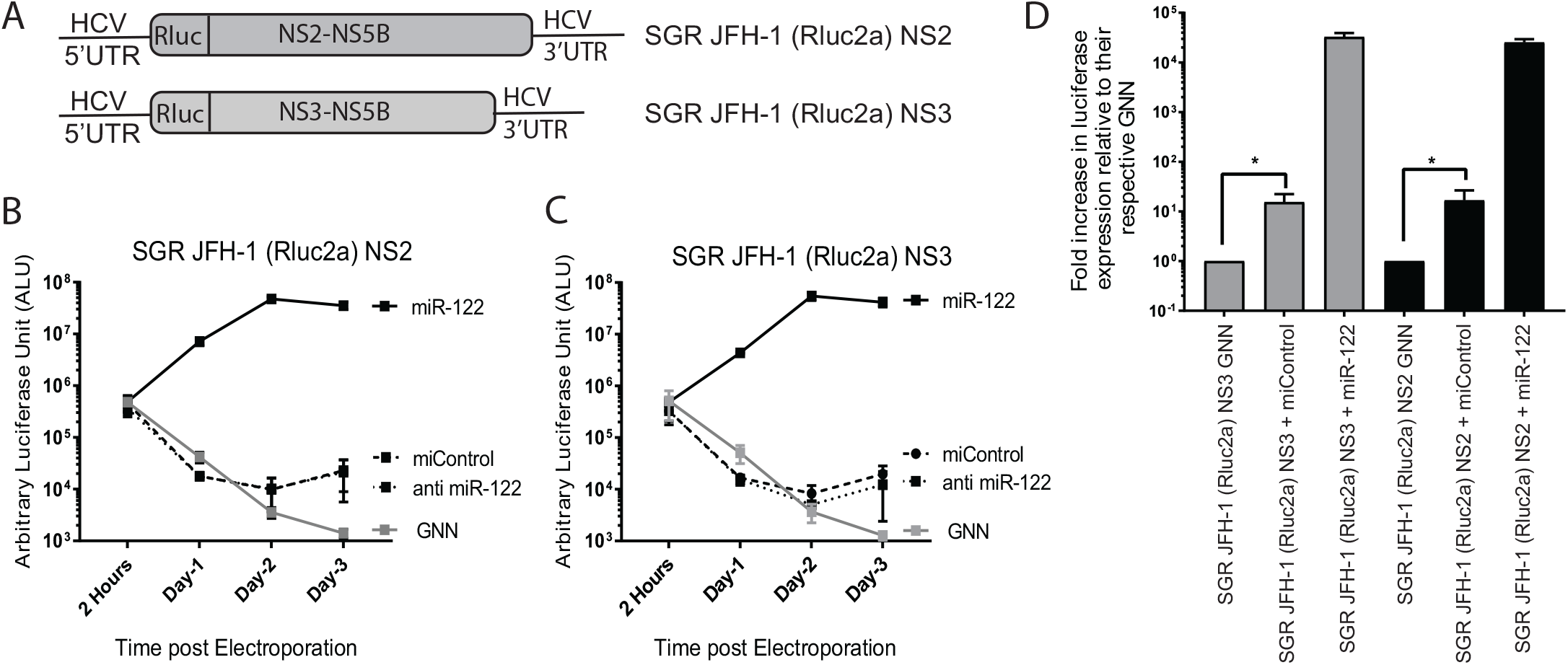
Deletion of the entire structural gene region facilitates miR-122-independent replication of HCV. (A) Schematic diagrams of the structural gene deletion mutants missing the entire structural gene region. SGR JFH-1 (Rluc2a) NS2 has a deletion of structural gene region and retaining either NS2 to NS5b non-structural genes, and SGR JFH-1 (Rluc2a) NS3 has a complete deletion of structural genes and NS2 and retaining NS3 to NS5b non-structural genes. Transient replication assay of (B) SGR JFH-1 (Rluc2a) NS2 and (C) SGR JFH-1 (Rluc2a) NS3 HCV RNA in the presence of a control miRNA (miControl) or miR-122 in Huh 7.5 miR-122 knock out cells, or anti-miR-122 LNA to verify miR-122-independent replication. GNN denotes replication assays using the respective replication defective mutants. (D) Fold increase in luciferase expression vs non-replicative GNN controls for SGR JFH-1 (Rluc2a) NS2 and (C) SGR JFH-1 (Rluc2a) NS3 HCV RNA adapted from data shown in (B) and (C). All data shown are the average of three or more independent experiments. Error bar indicates the standard deviation of the mean and asterisk indicates significant differences. The significance was determined by using Student’s t-test (ns-not significant, *P<0.033).

### HCV genomes capable of miR-122-independent replication replicate efficiently but in a small number of cells

Next, we wanted to characterize and compare cells supporting transient miR-122-dependent and miR-122-independent HCV replication. We hypothesized that low-level transient miR-122-independent HCV replication will be displayed as either a high number of cells that support low levels of HCV replication or alternatively, as a small number of cells that support efficient HCV replication. To test this hypothesis, we used immunofluorescence to detect the viral protein NS5a in infected cells and observed that a small number of cells support miR-122-independent replication compared to a high number of cells that support miR-122-dependent replication (**Fig. 5A**). We used flow cytometry to compare the intensity and number of cells stained with NS5a during miR-122-dependent *vs* miR-122-independent replication of HCV SGR JFH-1 Fluc RNA (∼2-6% and ∼45-60% respectively), U4C/G28A/C37U J6/JFH-1 (p7Rluc2a), a full-length viral RNA carrying three point mutations in the 5’ UTR previously reported to promote miR-122-independent replication of full-length genomic HCV RNA (∼2-6% *vs* ∼25-30%), (27), and J6/JFH-1 Rluc wild-type RNA a viral RNA that does not support miR-122-independent replication (∼0.2-1.5 *vs* ∼55-70%) (**Fig. 5 B, C**). In addition, the staining intensity appeared similar in individual cells supporting either miR-122-dependent or miR-122-independent replication (**Fig. 5A**), and the intensity of viral protein staining in NS5a-positive cells from miR-122-independent replication was equal to, or higher than in miR-122-dependent replication on day 3 post electroporation (**Fig. 5B, D)**. Thus, HCV RNAs capable of miR-122-independent HCV replication establish replication in a smaller proportion of cells than miR-122-dependent systems, but once established, viral protein levels appear similar. This suggests that miR-122 functions to enhance the establishment of HCV replication in a cell but appears dispensable to maintain efficient viral protein expression during the infection.

**FIG 5:**
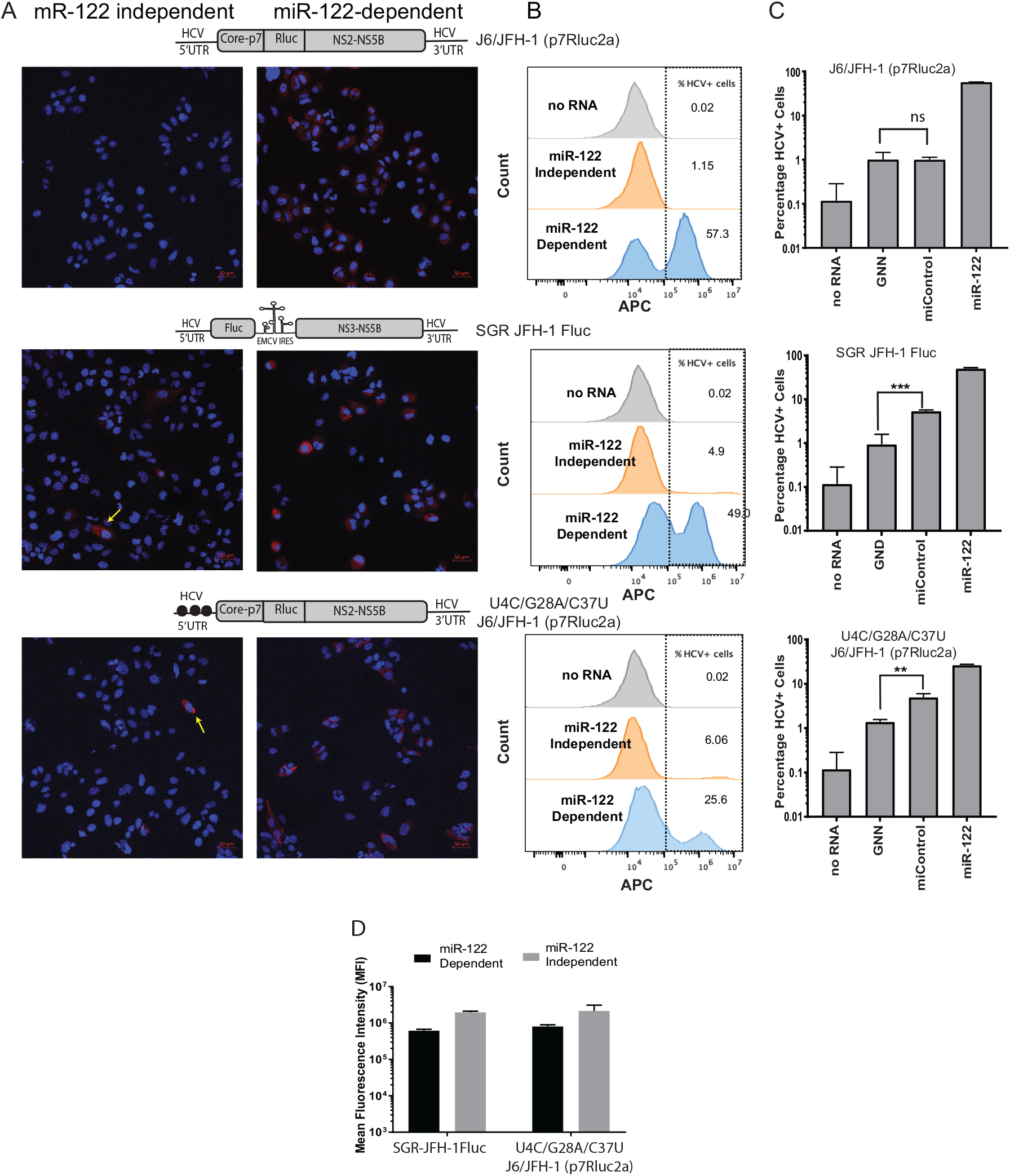
miR-122-independent replication is supported by a few cells within the population. (A) Immunostaining and confocal microscopy of Huh 7.5 miR-122 knock out cells co-electroporated with wild-type J6/JFH-1 (p7Rluc2a), SGR JFH-1 Fluc, and U4C/G28A/C37U J6/JFH-1 (p7Rluc2a) HCV RNA with control microRNA (miControl, Left Panel) or miR-122 (Right Panel). Cells were co-electroporated with viral RNA and microRNA and immunostained to detect NS5a on day 3. HCV NS5a is stained red with AlexaFluor-594 secondary and the cells were counterstained with DAPI to identify the nucleus. The respective RNA constructs are shown above the panel. Cells supporting miR-122-independent replication are indicated with yellow arrows. (B) Representative flow cytometry plots of Huh 7.5 miR-122 KO cells co-electroporated with J6/JFH-1 (p7Rluc2a), SGR JFH-1 Fluc, or U4C/G28A/C37U J6/JFH-1(p7Rluc2a) HCV RNA and control microRNA or miR-122. Cells were collected on day 3 post electroporation, stained for HCV NS5a with 9E10 anti-NS5a and APC-conjugated goat-anti mouse secondary antibody. The X-axis indicates the count of the cells and Y-axis indicates the intensity of the APC signal. Data are represented as histogram overlay of no RNA, miR-122-independent and miR-122-dependent replication. Grey histogram indicates the ‘no RNA’ control, whereas orange and blue histograms indicate the miR-122-independent and miR-122-dependent replication. No RNA control was used to gate HCV positive cells. (C) (D) (E) Percentage of Huh 7.5 miR-122 KO HCV positive cells electroporated with viral RNA [(C) wild-type J6/JFH-1 (p7Rluc2a), (D) SGR JFH-1 Fluc, (E) U4C/G28A/C37U J6/JFH-1 (p7Rluc2a)] and control microRNA or miR-122. Cells electroporated with no RNA, and GNN are used as negative controls. (F) The Mean Fluorescence Intensity of cells supporting HCV replication (Fig. B) was calculated with the FlowJo software ver 10.6. The MFI was measured as a proxy for viral NS5a protein accumulation in the cells. All data shown are the average of three or more independent experiments. Error bar indicates the standard deviation of the mean and asterisk indicates significant differences. The significance was determined by using Student’s t-test (**P<0.002, ***P<0.001).

### HCV variants supporting miR-independent replication do not show evidence of adaptation to growth without miR-122

An alternative explanation for efficient ongoing miR-122-independent replication of HCV is molecular adaptation of the viral genome to replication without miR-122. To identify potential adaptation to growth independent from miR-122 we used a biological assay that assessed the replication ability of viral RNA derived from cells supporting transient miR-122-independent replication. We used a biological method instead of sequencing as a rapid and sensitive way to detect if adaptation was a major influence in the observed miR-122-independent replication. We hypothesized that if viral RNA replicating in Huh 7.5 miR-122 KO cells had mutated to adapt to miR-122-independent replication, that the viral RNA derived from these cells would have an enhanced ability to replicate when electroporated into new Huh 7.5 miR-122 KO cells, and we would observe an increase in replication efficiency with repeated RNA passage through Huh 7.5 miR-122 KO cells. To test this, we harvested total cellular RNA from cells supporting miR-122-independent replication of SGR JFH-1 Fluc HCV RNA and electroporated 5 μg into of the total RNA into new Huh 7.5 miR-122 KO cells with and without miR-122 supplementation. miR-122-independent virus replication was measured based on Fluc expression in miR-122 KO cells electroporated with total RNA + miControl, and miR-122-dependent RNA replication was measured in miR-122 KO cells supplemented with total RNA + miR-122. The ratio of Rluc signal in miControl/miR-122 represented the relative efficiency of the viral RNA present in the total RNA sample to grow independent from miR-122 RNA (**Fig. 6A**). We also harvested and passaged RNA from cells supporting miR-122-dependent replication as a control. Based on miControl/miR-122 ratios over 3 RNA passages we did observe an increase in ratios between RNA passage 1 and passage 2 but the ratios did not differ from those of control RNA from cells harboring miR-122-dependent replication. In addition, miR-122-independent replication was undetectable in the total RNA after 2 RNA passages (**Fig. 6B**). Thus, we concluded that there was no detectible evolution of viral genomes with enhanced miR-122-independent replication.

**FIG 6:**
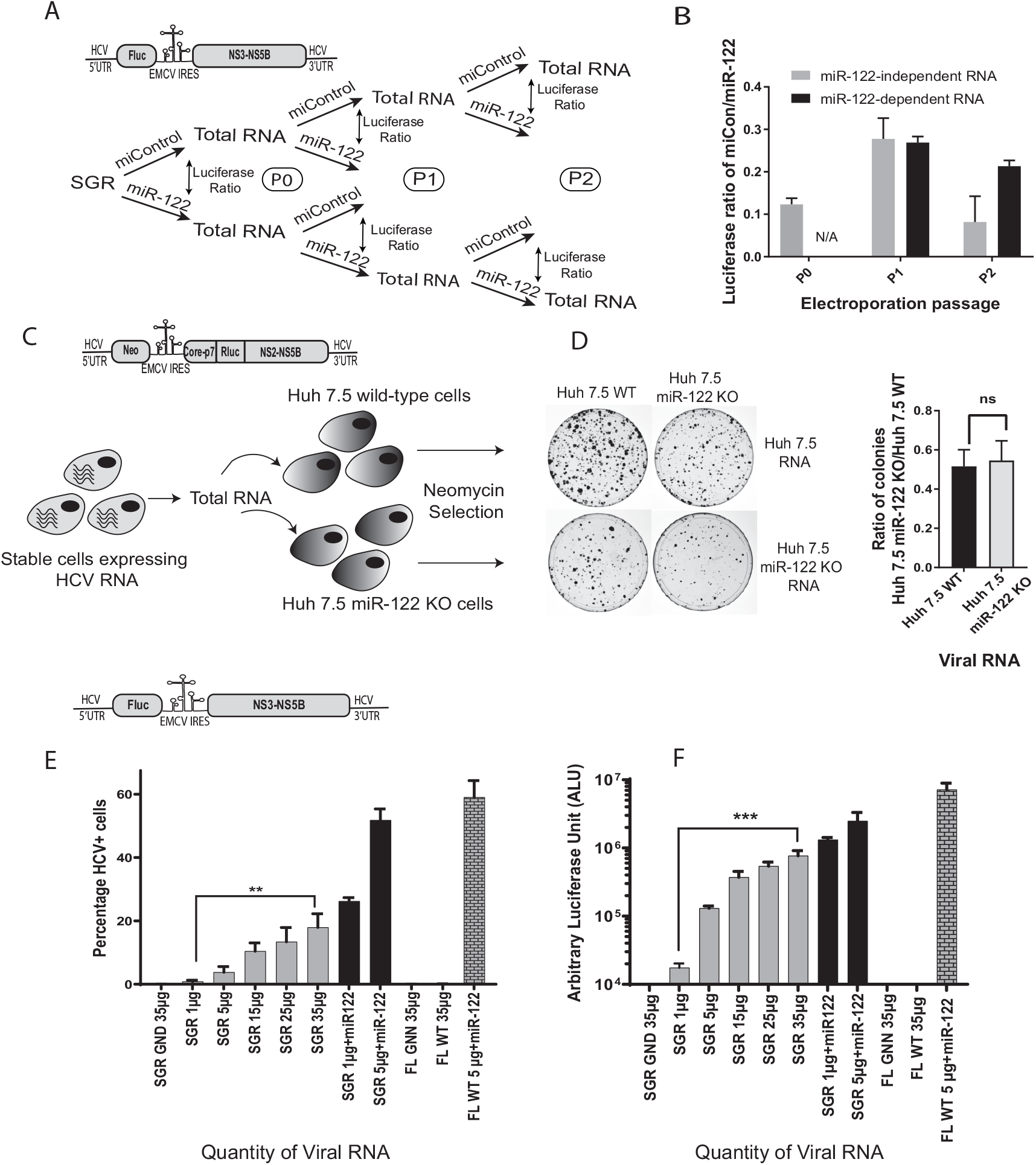
There was no evidence of genome adaptation to absence of miR-122 in HCV genomes passaged under miR-122-independent conditions. (A) Schematic diagram of the experiment to assess HCV adaptation to growth without miR-122. Total RNA was isolated from cells supporting transient miR-122-independent (miControl) or miR-122-dependent (miR-122) replication of HCV SGR JFH-1 Fluc and then re-electroporated into miR-122 KO Huh 7.5 cells with either miControl or miR-122. This was done for 3 RNA passages as shown. The relative luciferase expression when RNA was elecroporated with miControl was calculated vs the same RNA electroporated with miR-122 to determine the relative miR-122-independent replication ability of each RNA and plotted in (B). In (B) RNAs derived from miR-122 independent cells are noted as miR-122. (C) A schematic diagram of the experiment to test for HCV adaptation to in cells supporting stable miR-122-independent HCV replication. Huh 7.5 cells stably supporting miR-122-independent and miR-122-dependent replication of HCV J6/JFH-1 Neo (p7Rluc2a) were generated by electroporated of Huh 7.5 miR-122 KO or WT cells and selection with G418. Total RNA from Huh 7.5 cells stably supporting miR-122-independent or miR-122-dependent replication was isolated and electroporated into new WT or miR-122 KO Huh 7.5 cells and the ability to initiate miR-122-independent HCV replication was assessed based on colony formation and the ratio of colonies formed in Huh 7.5 miR-122 KO/WT cells (D). the constructs used for each experiment are shown above each diagram. (E and F) Transient replication assays showing the impact of the quantity of SGR JFH-1 Fluc RNA on the number of HCV + cells (E) and Fluc expression (F). In (E) the percentage of HCV positive cells was determined by using FACS and in (F) luciferase activity was assessed. SGR JFH-1 GND and J6/JFH-1 (p7Rluc2a) GNN (replication-defective mutants) were used as the negative controls. All data shown are the average of three or more independent experiments. Error bar indicates the standard deviation of the mean and asterisk indicates significant differences. The significance was determined by using Student’s t-test (ns-not significant, **P<0.002, ***P<0.001).

We next tested for HCV adaptation to miR-122-independent replication in cells stably supporting miR-122-independent replication. To test for adaptation, we assessed the ability of viral RNA extracted from Huh 7.5 miR-122 KO cells stably harboring J6/JFH-1 Neo Rluc RNA to form colonies when re-electroporated into naïve miR-122 KO cells. If adaptation had occurred, we expected a higher ratio of colonies formed after electroporation of total RNA derived from stable cells supporting miR-122-independent J6/JFH-1 Neo Rluc RNA replication compared with total RNA from cells supporting miR-122-dependent J6/JFH-1 Neo Rluc RNA replication (**Fig. 6C**). Similar to our transient adaptation assay, we did not observe a significant difference in the ratio of colonies formed following electroporation of total RNA derived from stable miR-122 KO vs stable miR-122 wild-type cell lines (**Fig. 6D**). Thus, we concluded that transient and stable miR-122-independent HCV replication was not due to observable adaptation of the viral RNA.

### The quantity of viral RNA affects the percentage of cells supporting miR-122-independent replication

Since we did not detect viral RNA adaptation to miR-122-independent replication, we wondered what mechanism led to some cells supporting efficient miR-122-independent replication and some cells remaining uninfected. Based on a role for miR-122 in stabilizing the HCV genome, we hypothesized that the cells supporting miR-122-independent replication were ones that had stochastically received sufficient viral RNA to initiate an infection. If this was the case then we surmised that electroporation of higher quantities of HCV RNA would increase the numbers of cells that received enough viral RNA to initiate an infection, and thus the numbers of cells supporting miR-122-independent replication should increase. Alternatively, if the cells supporting miR-122-independent replication had a specific phenotype that allowed for miR-122-independent HCV replication, then greater amounts of viral RNA would not enhance the numbers of cells supporting miR-122-independent replication. To test this, we electroporated increasing amounts of SGR JFH-1 Fluc HCV RNA into miR-122-knockout cells (5μg, 10μg, 15μg, 25μg, and 35μg) and measured the numbers of cells supporting miR-122-independent replication by flow cytometry. We observed an increase in the proportion of cells supporting HCV replication with an increased amount of viral RNA. A negative control sample electroporated with 35μg of non-replicative SGR JFH-1 Fluc GND RNA did not show any fluorescence signal. This data indicates that the amount of RNA affects the numbers of cells supporting miR-122-independent replication and suggests that the cells supporting miR-122-independent replication do not have a specific phenotype (**Fig. 6E**). Similar results were obtained in an experiment in which luciferase activity was used to assess viral replication and confirmed greater luciferase expression following higher amounts of virus RNA electropration (**Fig. 6F**). Further, this experiment also showed that using larger amounts of J6/JFH-1 WT viral RNA, which is incapable of miR-122-independent replication, did not enhance luciferase expression in miR-122 knockout cells. Thus, electroporation of higher amounts of viral RNA enhances miR-122-independent replication but did not enhance replication of wild-type genomes. This suggests that higher amounts of viral RNA can partially compensate for the lack of miR-122, and that the selection of cells that support miR-122-independent replication is likely stochastic. Since we have shown in our previous experiment that miR-122-independent HCV replication levels appears similar to miR-122-dependent replication, but in a fewer cells then we propose that cells supporting miR-122-independent replication had randomly received sufficient RNA to reach a threshold quantity required to initiate genome amplification. That electroporation of larger amounts of viral RNA can compensate for the lack of miR-122 is consistent with the proposed role for miR-122 in genome stabilization. However, because wild-type HCV RNA does not exhibit miR-122-independent replication, a high RNA concentration cannot compensate for the lack of miR-122. Thus it appears that genetic modifications that alter virus translation regulation such as an EMCV IRES or 5’UTR mutations are also necessary (35). This supports our and others’ findings that stabilization alone is not sufficient to promote miR-122 independent replication and both stabilization and altered translation regulation are required (14, 20, 36).

### miR-122 has a small but significant impact on the maintenance of an established HCV infection

Based on our data, miR-122 is required to establish an infection in a particular cell, but thus far appeared dispensable for ongoing maintenance of viral replication. To test this further we assayed the impact of miR-122 supplementation or depletion during later stages of the virus life-cycle. We first assessed the impact of miR-122 supplementation or antagonization on maintenance of transient miR-122-dependent HCV replication. For this assay we analyzed miR-122-dependent replication of J6/JFH-1 Rluc wild-type RNA, an RNA that cannot replicate without miR-122, in wild-type Huh 7.5 cells. J6/JFH-1 Rluc FL WT reached a maximum replication level at 3 days post electroporation and we supplemented or antagonized miR-122 at that time point and measured the impact on RNA replication by measuring Rluc expression two days later. After supplementing with miR-122, we saw a small but significant increase in luciferase expression from J6/JFH-1 Rluc FL WT **(Fig. 7A)**. Similarly, antagonization of miR-122 caused a small but significant decrease in luciferase expression **(Fig 7A)**. We observed similar results using ΔE2 J6/JFH-1 Rluc RNA, a mutant that cannot produce infectious particles and thus eliminates a possible impact of cell to cell spread of the infection. This suggests that miR-122 has a role, albeit small, in supporting ongoing miR-122-dependent HCV replication. Similarly, we tested the influence of miR-122 supplementation on the maintenance of miR-122-independent HCV replication. To test this, we used miR-122 KO cells supporting transient replication of a packaging-defective HCV mutant capable of miR-122-independent replication (U4C/G28A/C37U ΔE2 J6/JFH-1 Rluc HCV), and like in the previous experiment we supplemented the construct with miR-122 three days after viral RNA electroporation and measured the impact on Rluc expression two days later (**Fig. 7B**). In this case miR-122 supplementation did not change luciferase levels compared to miControl-supplemented cells during maintenance of replication. This data suggest that miR-122 does not have a significant additional impact on maintenance of viral RNAs capable of miR-122-independent replication after establishment of replication (**Fig. 7B**). Reports suggests that the mutations present on the 5’end of this viral RNA promote viral RNA stability (U4C), favours translation (G28A), and as well as promote viral genomic RNA synthesis by altering the structure of the negative strand (C37U) (19, 30, 35), and this may allow ongoing replication of this viral RNA in the absence of miR-122.

**FIG 7:**
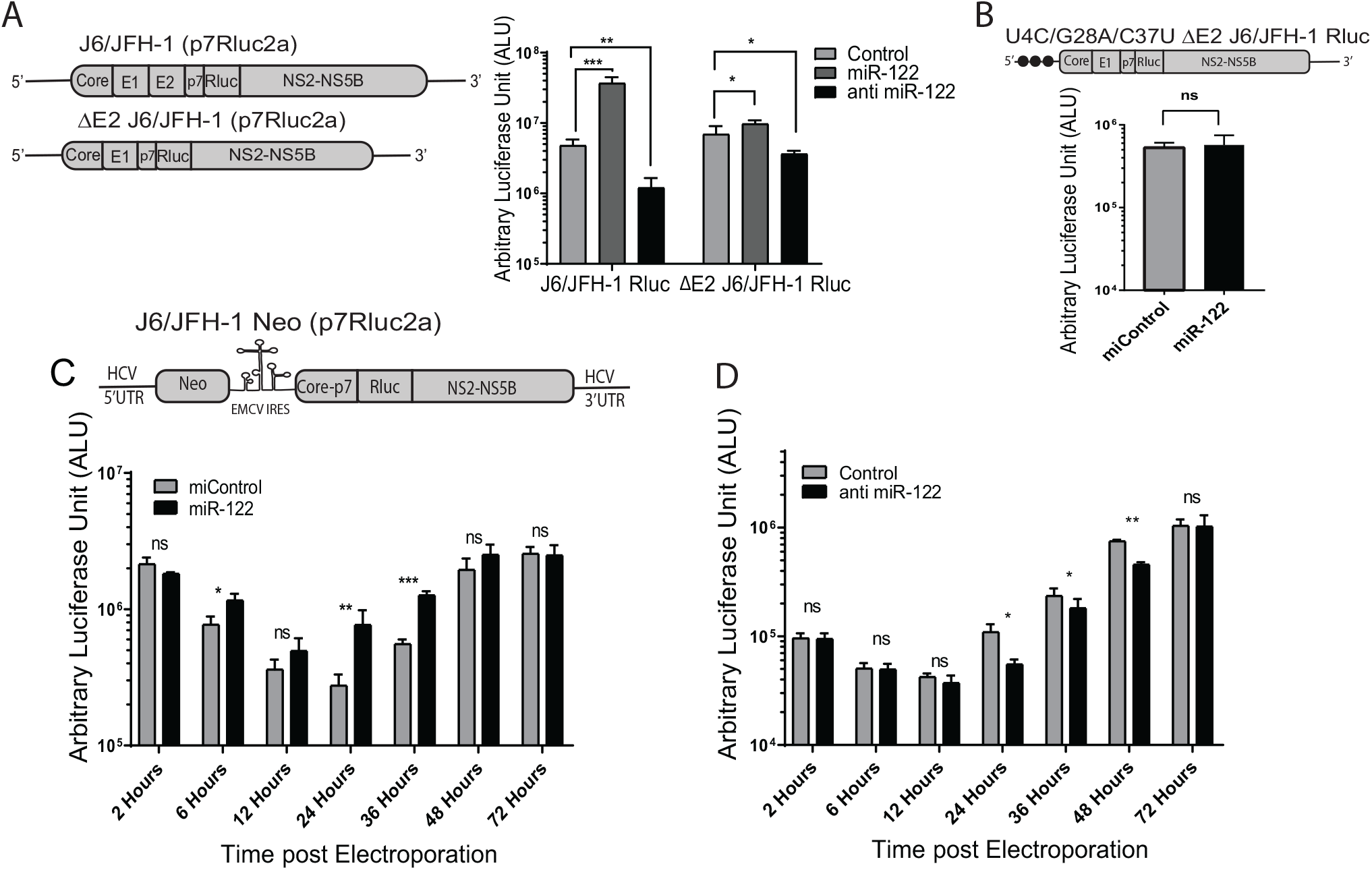
miR-122 has a small but significant influence on maintenance of HCV replication. (A) To test for the influence of miR-122 on ongoing HCV replication cells were supplemented with miR-122 or a miR-122 antagonist after the establishment of an infection. MiR-122-dependent replication of was established by electroporating Huh 7.5 miR-122 KO cells with wild-type J6/JFH-1 (p7Rluc2a) or ΔE2 J6/JFH-1 (p7Rluc2a) RNA and miR-122. 3 days later the cells were transfected with miR-122, anti-miR-122, or miControl to assess the influence of miR-122 supplementation, antagonization and 2 days later luciferase activity was measured as an indication of viral RNA amplification. (B) To assess the impact of miR-122 supplementation on HCV RNA replicating independently of miR-122, replication of U4C/G28A/C37U ΔE2 J6/JFH-1 (p7Rluc2a) HCV infection was established in Huh 7.5 miR-122 KO cells without miR-122 and on day 3, supplemented by transfection with either miR-122 or control microRNA. Luciferase activity was measured on day 2 post transfection as an indication of viral propagation. (C) To assess the impact of miR-122 supplementation on cells stably supporting miR-122-independent HCV RNA we used Huh 7.5 miR-122 KO cells electroporated with J6/JFH-1 Neo (p7Rluc2a) HCV and selected with G418. The cells were then electroporated with either control microRNA or miR-122, and luciferase activity was measured at different time points post-transfection. (D) To assess the impact of miR-122 antagonization on miR-122-dependent HCV replication we electroporated Huh 7.5 wild-type (WT) cells with J6/JFH-1 Neo (p7Rluc2a) HCV RNA and selected with G418. Stable cells were then transfected with either control LNA or anti-miR-122 LNA and luciferase activity was measured at different time points post transfection. All data shown are the average of three or more independent experiments. Error bar indicates the standard deviation of the mean and asterisk indicates significant differences versus the control microRNA/LNA. The significance was determined by using Student’s t-test (ns-not significant, *P<0.033, **P<0.002, ***P<0.001).

To further assessed the impact of miR-122 on ongoing HCV replication we also analyzed the impact of miR-122 supplementation and antagonization on cells supporting stable miR-122-dependend and -independent replication of J6/JFH-1 Neo Rluc HCV RNA (**Fig. 2A and E**). When cells supporting stable miR-122-independent replication were supplemented with miR-122, there was a small stimulation of luciferase expression observed at 24 and 36 hours post supplementation, but by 72 hours this effect was lost (**Fig. 7C**). In cells supporting miR-122-dependent HCV replication, electroporation of a miR-122 antagonist reduced luciferase expression at 24, 36, and 48 hours post electroporation, and again no difference was observed after 72 hours (**Fig. 7D**). These experiments suggest that miR-122 has a small but measurable influence on the maintenance of both miR-122-dependent and miR-122-independent HCV replication, but also suggests the influence is transient.

Altogether, our data show that miR-122 exerts its strongest effect by promoting the establishment of an infection in a cell by stimulating translation and stabilizes the viral genome and thus increases the chance that the virus will produce enough viral proteins to initiate an active infection before the RNA genome is degraded. In the absence of miR-122, the effect can be partially compensated for by using an alternative IRES to drive viral polyprotein translation or 5’ UTR mutations that increase the efficiency of viral protein translation (20, 27, 29) or by shortening the viral genome length, which may simply decrease the time to complete genome replication. Delivery of increasing amounts of viral RNA can also increase the chance of establishing miR-122-independent replication in a cell, likely by compensating for the function of miR-122 in genome stabilization. Finally, our data suggests a small, but significant positive influence on the maintenance of ongoing viral RNA replication. Based on these findings we suggest a model **(Fig. 8)** in which viral RNAs entering a cell are translated while being subject to degradation by host defence and RNA degradation pathways, but that miR-122 promotes translation and stabilizes the genome and thus increases the chances that enough viral proteins will be produced to initiate genome replication and establishment of virus replication complexes. In addition, we propose that miR-122 is required for ongoing HCV infections for similar reasons; to stabilize viral RNAs that exit the replication complex to initiate new replication complexes as the infection proceeds or when the infected cell divides.

**FIG. 8:**
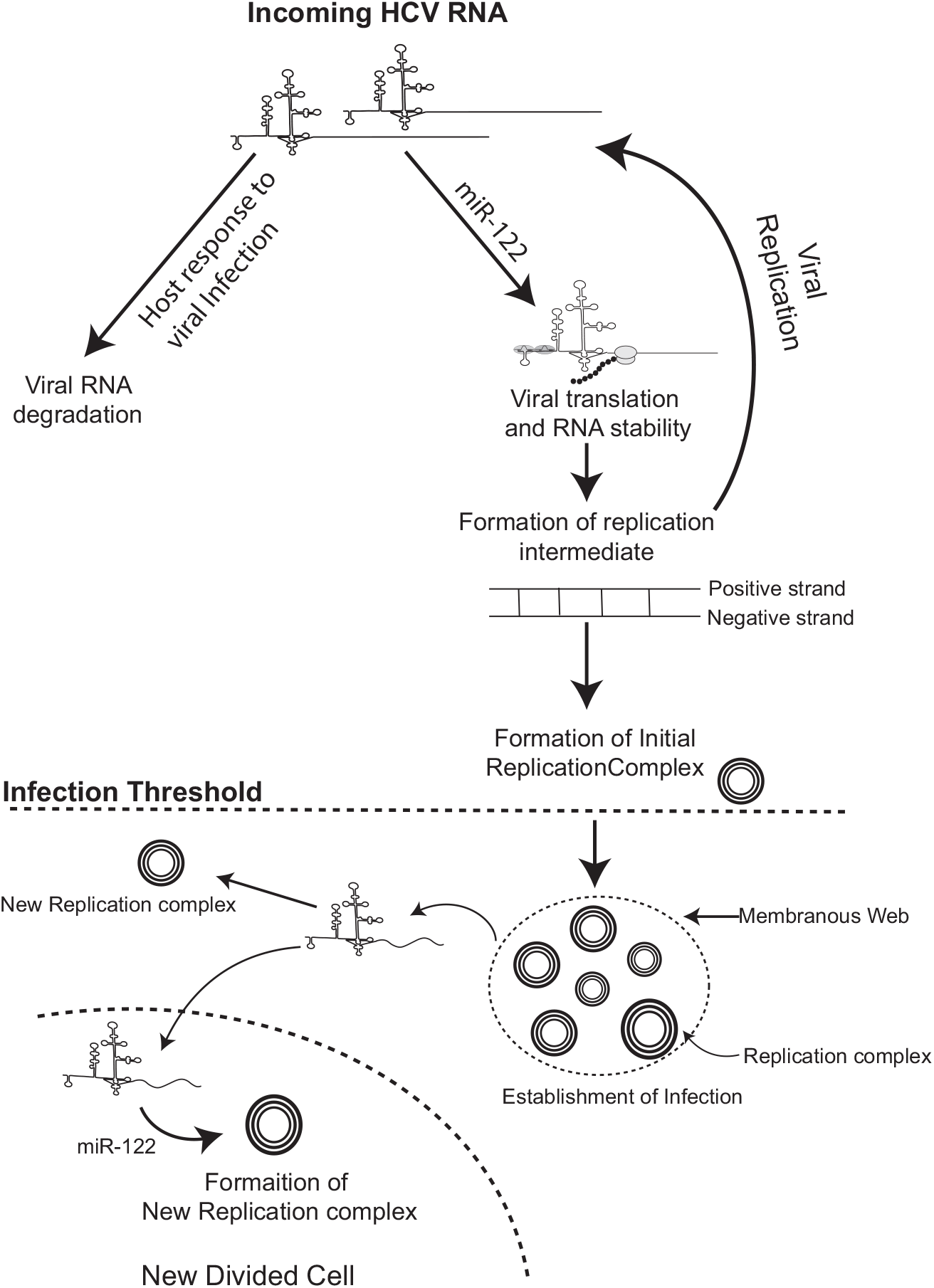
Proposed model of miR-122’s function in establishment of HCV infection. We propose that binding of miR-122 to the incoming viral RNA promotes viral translation and genome stability to allow the viral RNA to generate a threshold amount of viral protein required to initiate genome replication and the formation of virus replication complexes. After the establishment of replication complexes miR-122 may support ongoing virus replication by promoting the establishment of new replication complexes inside an infected cells or after an infected cell divides.

## DISCUSSION

While miR-122 is required for efficient and detectable HCV replication in cell culture, we and others have demonstrated that in certain contexts HCV can replicate independent from miR-122 (20, 25–27, 29, 37). Previously, we have identified that bicistronic HCV subgenomic replicons, and HCV genomes with point mutations within the miR-122 binding region of the 5’ UTR promote miR-122-independent replication (17, 20, 26). In this paper, we have shown that miR-122-independent replication is facilitated by the addition of an EMCV IRES (**Fig. 2**) and by a smaller viral genome with the deletion of structural genes (**Fig 3**). Since viral IRES elements regulate cap-independent translation, we believe that the presence of the EMCV IRES enhances the overall translation efficiency of the viral polyprotein and thus substitutes for the function of miR-122 in stimulating translation. However, the efficiency of miR-122-independent HCV replication is significantly lower than the miR-122-dependent replication, and thus the presence of an EMCV IRES cannot completely rescue the viral replication in the absence of miR-122. Our data have also suggested that shorter viral genomes with the deletion of whole structural gene region can replicate to low but detectable levels in the absence of miR-122 (**Fig 4**), even in viral genomes that lack an EMCV IRES or 5’ UTR point mutations. We speculate that the small genome size resulting from the removal of the structural gene region allows for more rapid virus protein translation and RNA replication simply based on the size of the protein product and genome, but we cannot omit the possibility that the structural gene region somehow inhibits viral propagation in the absence of miR-122. Thus, 5’ UTR point mutations, an EMCV IRES, and removal of the structural protein region each facilitate detectable HCV replication in the absence of miR-122.

We also observed that miR-122-independent replication is manifested as efficient genome replication within a small percentage of cells of a population, rather than inefficient genome replication in a large proportion of the cells (**Fig 5**). This suggests that miR-122-independent RNA replication is efficient but is initiated in only a small number of cells. We propose that the few cells that support miR-122-independent replication stochastically received enough RNA to initiate an infection and do not have a cellular phenotype that promotes initiation, as evidenced by increasing amounts of viral RNA resulting in increasing numbers of cells supporting miR-122-independent replication (**Fig. 6 E and F**). In this context, we propose that the EMCV IRES and the small genome size facilitate genome translation and genome replication such that the threshold for establishment of an active HCV replication complexes is lower. These findings also support the notion that miR-122 promotes the virus life-cycle by stabilizing the genome and stimulating virus translation, influencing the ability of the virus to establish an infection within a particular cell. Sudies on HCV capable of replicating independent of miR-122 showed that enhanced viral translation and stability can completely rescue HCV replication in the absence of miR-122 (35, 36), further confirming miR-122’s role in viral translation and genome stability in early stage of viral infection. Thus, we propose that miR-122 stabilizes the genome to provide sufficient viral genomes and protein translation to jump-start the early stages of an infection.

Microscopy and flow cytometry data also suggested that once the infection is established in the absence of miR-122, viral protein levels are equivalent to infection established in the presence of miR-122. This finding supports the data published by the Matsuura group that miR-122 is essential for the initial stage of viral infection but are dispensable for maintenance of infection (25). In addition, Villanueva *et al*. showed that the addition of miR-122 or anti-miR-122 has no effect on HCV RNA synthesis *ex vivo* in membrane-bound replicase complexes isolated from HCV infected cells. They also observed no significant detectable quantities of miR-122 associated with replicase complexes *in vivo*, suggesting no significant role of miR-122 on the elongation phase of viral RNA synthesis (38). By contrast, we have found that miR-122 has a small effect in maintenance of viral replication. This supports work published by Masaki *et al*. (18) who proposed a direct role for miR-122 in promoting genome amplification. However, based on our data, we propose an alternative hypothetical model that miR-122 promotes the establishment of new replication complexes within an already-infected cell; an extension of its effect at infection initiation. This is supported by our experiments showing a requirement for miR-122 from 24 to 36 hours post miR-122-supplementation in stably infected cells, but not after 72 hours (**Fig. 7**). We speculate that this is the time when there is active cell growth, and by 72 hours the cells have become confluent and we propose that miR-122 enhances the rate at which new replication complexes form as infected cells divide. This could be through a direct role of miR-122 in promoting genome amplification as proposed by Masaki *et al*., but is also consistent with the roles for miR-122 in stabilizing and stimulating translation of newly synthesized viral RNAs as they are released from one replication complex to promote their initiation of new replication complexes. Indeed, miRNAs and Ago have been implicated in liquid-liquid phase separation and sequestration of mRNAs into protein complexes, and miR-122 could also function to directly stimulate replication complex formation (39). This would explain why Vilanueva *et al*. did not see any effect of miR-122 supplementation or antagonization on HCV propagation in isolated membrane-bound replicase complexes, as they were assayed *ex vivo* without provision for the formation of new replication complexes. Interestingly, however, miR-122 supplementation did not affect the ongoing replication of the mutant HCV (U4C/G28A/C37U) capable of replicating independent of miR-122. A recent report suggests that these mutations substitute for miR-122 by stabilizing the viral genome (U4C), promoting viral translation (G28) and promoting the synthesis of positive strand RNA (C37U) (30) and this could explain why miR-122 supplementation had no influence on the ongoing replication of the virus. Overall, our model **(Fig. 8)** posits that miR-122 promotes genome stabilization and translation to initiate replication complex formation to initiate an infection but may similarly affects newly synthesized viral RNAs to promote the development of new replication complexes within an infected cell and in daughter cells following cell division.

Initiation of an HCV infection in the liver is likely more dependent on miR-122 than in tissue culture cells since the HCV RNA copy number varies from less than 1 to 8 per hepatocyte in the *in vivo* condition (40), 100 fold lower than in tissue culture cells. Thus, despite the abundance of miR-122 in hepatocytes, infection with a single or a low copy number of viral genomes may requires multiple rounds of viral replication to establish an infection and could explain the importance of miR-122 in both the establishment and ongoing viral replication in the liver. In addition, recent findings reported that other RNA viruses such as enterovirus, norovirus, and rotavirus transmit among hosts in vescicle cloaked viral clusters that are found to be more infectious than the single free viral particles (41, 42). Similarly, miR-122 may also enhance the viral genome quantity by slowing RNA degradation. Further, although there are no reports of HCV virion clusters, exosomes containing replication competent HCV RNAs in complex with miR-122, Ago-2 and HSP90 can transmit HCV infection which could increase the copy number and stabilty of infecting HCV genomes (43).

The use of anti-miR-122 LNA (Miravirsen, SPC3649) has shown promising results as a therapeutic agent, as its use in infected patients decreased HCV levels to undetectable levels (44– 46). Our findings suggest that anti-miR-122 exerts its effect primarily during the establishment of HCV infection and suggest that infections in the liver must be highly dynamic such that new cells are continually being infected, and antagonization of miR-122 might be restricting new cell infection and decrease the overall viral burden of the liver. This also aligns with the hypothesis proposed by Stiffler *et al*. which suggests that HCV infection might be a transient infection *in vivo*, and the chronic condition of the virus is maintained by a continuous cycle of viral infection and clearance. However, the low HCV genome copy number in infected cells *in vivo* could also make HCV infections more reliant on the supportive role of miR-122 during ongoing infections thus miR-122 antagonism may also decrease HCV replication levels within infected liver cells.

Since Miravirsen targets a host factor it has a high barrier to resistance and is effective against all HCV genotypes (44, 47). However, our work and that of others caution that HCV can replicate independent from miR-122 and can develop resistance to miR-122 targeting therapy. Resistance-associated substitutions (RAS) variants G28A (Guanine is replaced by Adenine at position 28 of viral genome) and C3U (Cytosine is replaced by Uracil at position 3 of viral genome) were identified in a patient treated with Miravirsen, and are capable of replicating at a low abundance in the absence of miR-122 (23, 48). In addition, several studies have identified mutants capable of replicating independently of miR-122 in cell culture (20, 27). Thus, the application of anti-miR-122 should be done with care. The addition of Miravirsen to the DAA treatment has been shown to be synergistic in suppressing HCV replication and thus rather than being a stand-alone therapeutic, Miravirsen may instead be best used as a part of the treatment cocktail for patients who are non-responsive to first-line DAAs (49).

Overall, in this study we show that the influence of miR-122 can be partially compensated for by using an alternative IRES elements to drive viral polyprotein translation, 5’ UTR mutations that increase the efficiency of viral protein translation or by shortening the viral genome length, and we speculate that this increases the efficiency by which the viral genome can be translated and replicated. Next, we showed that miR-122-independent replication is displayed as efficient replication in a smaller number of cells compared to miR-122-dependent replication. This suggested that miR-122 is required to initiate an infection in each cell but we have found that miR-122 does have a small but measurable impact on ongoing replication. Based on these findings we propose a model **(Fig. 8)** in which miR-122 stabilizes the viral genome and stimulates translation, thus increases the chance that the virus will produce sufficient viral proteins to initiate an active infection before the RNA genome is degraded. In addition, we propose that miR-122 is required for ongoing HCV infections for similar reasons; to promote the generation of new replication complexes as the infection proceeds or as the infected cell divides. Thus, we propose that a successful viral infection is an interplay between viral protein synthesis, RNA replication, and RNA degradation, and that miR-122 favors the successful formation of the replication complex and the establishment of an infection.

## MATERIALS AND METHODS

### Viral cDNA Plasmids

Plasmids pSGR JFH-1 Fluc WT and pSGR JFH-1 Fluc GND contain bicistronic JFH-1-derived subgenomic replicon (SGR) cDNAs with a firefly luciferase reporter and an Encephalomyocarditis Virus (EMCV) IRES, and were provided by Dr. T. Wakita (50). pSGR GND is the control nonreplicative version containing a mutation to the viral polymerase active site, GDD, to an inactive GND. Plasmids encodng a full-length HCV genome expressing a *Renilla* luciferase (Rluc) gene, pJ6/JFH-1 RLuc (p7RLuc2A) (51), and a full-length bicistronic HCV cDNA expressing neomycin from the HCV IRES and Rluc within the full-length HCV polyprotein, pFLneo-J6/JFH-1 (p7-Rluc2a) were provided by Dr. C. M. Rice (herein called pJ6/JFH-1 Rluc and pJ6/JFH-1 Neo Rluc respectively) (51). A GDD to GNN mutation rendering the viral polymerase nonfunctional in the negative control pJ6/JFH-1 Rluc GNN was also provided by Dr. C. M. Rice (51). Plasmids pFGR Fluc JFH-1 WT and pFGR Fluc JFH-1 GNN bear full-genomic bicistronic replicons of JFH-1, and a firefly luciferase reporter (52). Plasmid pJ6/JFH1 Rluc encoding triple mutations in the 5’UTR of the virus genome (U4C/G28A/C37U) was made by site-directed mutagenesis of a smaller plasmid (pBSKS+) encoding *EcoR*I to *Kpn*I fragment of J6/JFH-1 Rluc. Once the mutations were confirmed in pBSKS+, the *EcoR*I to *Kpn*I fragment from pBSKS+ was cloned into pJ6/JFH-1 to obtain U4C/G28A/C37U pJ6/JFH-1 Rluc. HCV structural gene deletion mutants were constructed in pJ6/JFH-1 Rluc plasmid using 5’Phosphorylated primers to perform inverse PCR. Unmodified template was then removed by DpnI digestion, and PCR products were then cirularized with T4 DNA ligase. All the deletion mutants were sequenced and confirmed for deletions.

ΔCore: 5’Phos-CCTTTTTCTTTGAGGTTTAGGATTTG3’-F 5’

Phos-TGCTCCTTTTCTATCTTCTTG3’-R

ΔE1: 5’Phos-GGAGCAACCGGGTAAGTTCC3’-F 5’

Phos-AAAGTCGTTGTCATCCTTCTG3’-R

ΔE2: 5’Phos-CTTCTGTTGGCCGCCGGGGTGGAC3’-F 5’

Phos-CAGGCCGAAGCAGCACTAGAGAAGC3’-R

ΔE1E2: 5’Phos-GCATTGCCCCAACAGGCTTATGC3’-F 5’

Phos-GCCGGTACTGATGTTCTTCACTTC3’-R

J6/JFH-1 Rluc with deletion of envelope protein 2 (ΔE2) was used to construct U4C/G28A/C37U ΔE2 pJ6/JFH-1 Rluc HCV where *EcoR*I to *Age*I fragment of ΔE2 pJ6/JFH-1 Rluc plasmid was replaced with *EcoR*I to *Age*I fragment from U4C/G28A/C37U pJ6/JFH-1 Rluc. pΔE1E2 J6/JFH-1 Rluc Plasmid was obtained from Dr. Ralf Bartenschlager (52).

### Construction of monocistronic and bicistronic sub-genomic constructs

Monocistronic pSGR JFH-1Rluc NS2 was made from J6/JFH-1 (p7Rluc2a) by generating a short intermediate PCR product using primers SGR NS2 F and R1, and then using this product as a forward megaprimer in tandem with SGR NS2-R2 the region from core to p7 was removed.

SGR NS2: 5’CGACGGCCAGTGAATTCTAATAC3’-F

5’CTGGATCATAAACTTTCGAAGTCATAGGCCGGCCGGTTTTTCTTTGAGGTTTAGGA TTTG3’-R1

5’CGGCCCATATGATGCCATCG3’-R2

Once confirmed through sequencing, the *EcoR*I to *Not*I region of the plasmid was swapped in to the original J6/JFH-1 (p7Rluc2a) plasmid to create pSGR JFH-1Rluc NS2. pSGR JFH-1Rluc NS3 was generated with the same approach using primes SGR NS3 F, R1 and R2 (Supplementary **Table S1**) to remove the NS2 region from the pSGR J6/JFH-1Rluc NS2, and then the *EcoR*I to *Spe*I fragment was inserted into the original JFH-1 Rluc to create pSGR JFH-1Rluc NS3.

SGR NS3: 5’GCCTCGTGAAATCCCGTTAG3’-F

5’GCTGGGCATAAGCAGTGATGGGAGCGGGCCCTGGGTTGGACTCGACGTC3’-R1

5’TCCACACTTGCACGGCTCCAAAGAC3’-R2

Replication -defective mutant controls were generated by cloning the GNN fragment from pJ6/JFH-1 (p7RLuc2a) GNN with *SnaB*I and *Xba*I into the monocistronic and bicistronic HCV plasmids. All modifications were confirmed by sequencing.

### *In vitro* RNA transcription

Viral RNAs were transcribed in vitro using the MEGAScript T7 High Yield *in vitro* Transcription Kit (Life Technologies, Burlington, ON, Canada). HCV cDNA plasmids were linearized with *Xba*I and made blunt with mung bean nuclease and transcribed as suggested in our previous studies (17).

### Cell Culture

Huh 7.5 (WT) cells, Huh 7.5 miR-122 KO cells, and Huh 7.5 cells stably expressing J6/JFH-1 Neo Rluc were grown and maintained as described by Amador-Cañizares *et al*. (20).

### Transient HCV replication assays

6.0×10^6^ cells suspended in 400 μl 1xPBS were co-electroporated with 5μg of HCV genomic RNA (unless otherwise indicated) and 3uL miR-122/miControl/anti-miR-122/anti-miR-124 at 10mM concentration and resuspended in 4 ml cell culture media (DMEM + 10% FBS + 1nM non-essential aminoacid + 100 μg/ml Pen/Strep). A total of 500 μL of the electroporated cells were plated onto each well of a 6-well plate in culture medium and incubated at 37^0^C.for the indicated lengths of time.

### Analysis of the effects of miR-122 supplementation and antagonization on transient HCV replication

To analyze the impact of miR-122 supplementation or antagonization on transient HCV replication assays, cells were co-electroporated with HCV RNA and miR-122 as described above, and a total of 250μL of re-suspended cells were allowed to grow on 6 well plates for 3 days. On day 3 the cells were transfected with 60 pMol of either miR-122 or control microRNA (miR-122 p2-8) or anti-miR-122 LNA or control LNA (anti-miR-124) with 5uL of Lipofectamine 2000 (Invitrogen). Cells were harvested 2 days post-transfection and luciferase activity was measured.

### Establishment of miR-122 KO stable cell line expressing bicistronic pJ6/JFH-1 Neo Rluc

Huh 7.5 miR-122 KO cells were electroporated with 10μg of pJ6/JFH-1 Neo Rluc RNA and all the cells were plated on a 15 cm tissue culture plate with DMEM+800 μg/ml G418 sulfate. Cells were selected for more than 60 days and flow cytometry was used to compare the cells with the already-established Huh 7.5 cells stably expressing pJ6/JFH-1 Neo Rluc (data not shown).

### Analysis for adaptation to replication without miR-122 during transient replication assays

Initially, 5 μg of SGR (JFH-1) was co-electroporated with miR-122 or miControl into 6.0×10^6^ Huh 7.5 miR-122 KO cells, and then resuspended into 4 ml of cell culture media. 500μL of cells are plated in 6 well plates for luciferase as well as for total RNA. Total RNA was extracted at each passage and a total of 10μg of total RNA was again co-electroporated with either miR-122 or miControl and the process was repeated for 3 passages.

### Analysis for adaptation to replication without miR-122 in stable HCV cells

Total RNA was isolated from Huh 7.5/ Huh 7.5 miR-122 KO stable cell lines and 10 μg of total RNA was electroporated into approx. 6.0×10^6^ Huh 7.5 and Huh 7.5 miR-122 KO cells. After electroporation cells were resuspended into 4 ml of cell culture media. A total of 1 or 2 ml of cells were plated on a 10 cm tissue culture dish containing DMEM+ 800 μg/mL G418 sulfate and incubated at 37 for approximately 2 weeks till the colonies develop. The plates were then washed with1x PBS, fixed with ice-cold methanol, and stained with crystal violet. Plates were scanned and colony numbers were quantified using ImageJ.

### Analysis of the effects of miR-122 supplementation and antagonization on stable HCV replication

Huh 7.5 WT/Huh 7.5 miR-122 KO cells expressing pJ6/JFH-1 Neo Rluc were electroporated with miR-122 or control microRNA. Cells were plated on a 6-well plate and harvested at different time points according to above-mentioned protocol for luciferase analysis. For transfection, approximately 5.0×10^5^ cells were plated a 6 well plate. 60 pmol of miR-122/control microRNA was transfected to the cells next day using 5 μL lipofectamine 2000. Cells were harvested 48 hours post transfection and luciferase activity was measured.

### Luciferase assays

Cells were washed with 1X Dulbecco’s PBS and harvested with 100 μL of 1X passive lysis buffer diluted in ddH_2_O (Promega, Madison WI, USA). Luciferase activity in cell lysates were measured by using either Firefly or *Renilla* luciferase kits (Promega, Madison WI, USA), and light emission was measured in arbitrary light units on a Glomax 20/20 Luminometer (Promega, Madison WI, USA). The luciferase assays were performed as suggested by the manufacturer’s protocols.

### Total cellular RNA extraction

1 ml of Trizol was used to harvest the total cellular RNA, following the suggested manufacturer’s protocol (Life Technologies, Burlington, ON, Canada).

### Immunostaining for Confocal Microscopy

150 μL of electroporated cells were placed on (18 mM diameter, 1.5 thickness) coverslip and placed in a12 well plate with culture media. On day 3 cells were washed with PBS and fixed with 4% paraformaldehyde. After washing three times with PBS, cells were permeabilized and blocked with 0.1-0.2% Triton X 100 + 3% BSA in PBS for 1 hour. Mouse anti-NS5a 1°Ab (9E10, provided by Dr. Charles Rice (53)) was incubated on the cells at 1:5000 dilution in the blocking buffer overnight at 4°C. Cells were further washed three times with washing buffer (0.02-0.01% Triton X-100 + 0.3% BSA in PBS) and incubated with 2° Ab at 1:1000 dilution (Goat Ab-AF 594, Life Technologies, A11005) for 1 hour at room temperature. After washing the cells three times with washing buffer, 1 μg/ml DAPI in PBS was used to stain the nucleus for 10 minutes. Cells were further washed with PBS and ddH_2_O to remove the residual salts and the coverslip was mounted on a glass slide using SlowFade Gold antifade mounting solution (Life Technologies). Images were captured using a Zeiss confocal microscope (LSM 700) and images were processed using Zen 3.1 (Blue edition) and ImageJ software.

### Immunostaining for Flow cytometry

On Day 3 of electroporation cells were harvested from 6-well plates and flow was done as described by Kannan *et al*. (54). Primary Ab against HCV NS5a was used at a dilution of 1: 2500 (9E10, Mouse anti NS5a), and Secondary Antibody APC: (F ab’) 2 Goat Anti-Mouse IgG APC (eBioscience, Cat 17-4010-82) was used as suggested by the manufacture. Samples were suspended in 300 μL of 1XPBS and acquired in Beckman Coulter CytoFlex flow cytometer. Data were analyzed by FlowJo ver10.6

### Statistical analysis

All data are displayed as the mean of three or more independent experiments, and error bars indicate the standard deviation. Statistical analysis was performed using Graph Pad Prism v7, and the statistical tests are indicated in the figures.

## ACKNOWLEDGMENTS

We would like to acknowledge Dr. Charlie Rice (The Rockefeller University) for providing the J6/JFH-1(p7Rluc2A), FL-Neo J6/JFH-1(p7Rluc2A) and Huh-7.5 wild-type cells. We also acknowledge Dr. Matthew Evans (Icahn School of Medicine at Mount Sinai) for Huh-7.5 miR-122 knockout cells. We would also like to thank Saurav Saswat Rout for his help with flow cytometry data analysis.

## AUTHOR’S CONTRIBUTIONS

M.P, P.A.T, and J.A.W. designed and performed the experiments, and analyzed the data; M.P and J.A.W. wrote the manuscript.

